# Hierarchical unimodal processing within the primary somatosensory cortex during a bimodal detection task

**DOI:** 10.1101/2022.08.12.503802

**Authors:** Sergio Parra, Héctor Diaz, Antonio Zainos, Manuel Alvarez, Jerónimo Zizumbo, Sebastián Pujalte, Lucas Bayones, Ranulfo Romo, Román Rossi-Pool

## Abstract

Where and how in the brain do neurons process more than one sensory modality? To answer these questions, scientists have generated a wide variety of studies at distinct space-time scales in different animal models, and often shown contradictory conclusions. Some conclude that this process occurs in early sensory cortices, but others that this occurs in areas central to sensory cortices. Here, we sought to determine whether sensory neurons process and encode physical stimulus properties of different modalities (tactile and acoustic). For this, we designed a bimodal detection task where the senses of touch and hearing compete from trial to trial. Two Rhesus monkeys performed this novel task, while neural activity was recorded in areas 3b and 1 of the primary somatosensory cortex (S1). We analyzed neurons’ coding properties and variability, organizing them by their receptive field’s position relative to the stimulation zone. Our results indicate that neurons of areas 3b and 1 are unimodal, encoding only the tactile modality, both in the firing rate and variability, but not to the acoustic one. Moreover, we found that neurons of both subareas encode the tactile information differently, revealing a hidden processing-based hierarchy. Finally, using a powerful non-linear dimensionality reduction algorithm, we show that the activity from areas 3b and 1 can be separated, establishing a clear division in the functionality of these two subareas of S1.

**SIGNIFICANCE STATEMENT:** Our brain integrates information from all our senses to perceive the external world. But where and how in the brain this integration occurs? Here we ask if the primary somatosensory cortex (S1) encodes information from more than one sensory modality. We recorded the activity of single neurons from areas 3b and S1, while trained monkeys performed a bimodal detection task, where tactile and acoustic stimuli compete. The analysis showed that neurons from areas 3b and 1 responded only to the tactile modality both in their rate and variability. However, our results support that these two areas are different enough as to be considered functionally distinct entities.

## INTRODUCTION

Have you ever turned down the volume on your car’s radio when looking for a place to park or closed your eyes when trying to detect a sound? Sometimes, the inputs from our different senses can interfere or compete with each other, even though integrating them is essential to form a coherent perception of the external world. Neural correlates of this competition should emerge in areas of the brain that participate in multisensory integration. The search for where and how in the brain this operation is carried out is an active area of research. In particular, the question of whether primary sensory cortices have any explicitly multisensory responses has been the subject of much discussion. The mainstream view suggests that this is not the case; primary sensory cortices are unimodal, processing only their own sensory modality. In this work we investigate if areas 3b and 1 of the primary somatosensory cortex (S1) show multisensory processing. We do so by analyzing the firing rate coding and variability of single neurons, recorded while Rhesus monkeys performed a bimodal detection task (BDT) that involved competition between the senses of touch and hearing.

It was thought that multisensory processing, at the neural level, came only after intensive unimodal processing (1, 2). Surprisingly, recent decades have brought several studies arguing that it also occurs in areas considered thoroughly unimodal, such as the primary sensory cortices (3–13). This would mean that S1, for example, not only processes tactile but also auditory and even visual information (3, 14). Similarly, the auditory cortex would process tactile and visual stimulus properties in addition to auditory ones. However, this growing body of evidence comes mainly from studies that use recording techniques with low (EEG, fMRI) or medium (LFP) spatial resolution, but even in single unit recordings (3, 10, 14). Further, their experimental designs have led to inconclusive findings due to lax control of stimulus properties or the simultaneous presentation of complex stimuli from different modalities. This could confound multisensory responses with unimodal ones to uncontrolled properties that are correlated between stimuli of different modalities.

For our study and assuming the unimodal tactile processing for S1, we asked two core questions for the acoustic modality: Is there a change in the firing rate of S1 neurons during an acoustic stimulus? In their variability? Is there any difference in the response properties of neurons in 3b vs area 1 during acoustic stimulation? There is, nonetheless, an alternative hypothesis to multisensory processing: attention, which could explain some of the differences in neural responses between unimodal and multimodal tasks. However, previous evidence (15) has shown that performance in detection tasks is not affected by dividing attention between two senses. Consequently, we focus on bimodal detection in our present study. Still, we do comment further on attention in the Discussion.

This work contributes evidence that could address some of the drawbacks of previous studies. First, electrophysiological data were recorded with high spatial resolution. We obtained spike trains from single neurons using independently moving microelectrodes and a custom offline spike sorting algorithm. Second, a bimodal detection task (BDT) was used, in which, interleaved vibrotactile and acoustic stimuli of varying amplitude were delivered. This means that the stimuli presented could be at or below the threshold for detection, putting more emphasis on the responses of S1 (16). Third, we analyzed both the stimulus coding and variability of spike trains. On the one hand, coding could be considered the foremost functional feature of neural responses for cognition, and thus is fundamental for understanding the role of an area in the brain processing. On the other, variability is an important metric in which coding is constrained and has rarely been studied in the context of multisensory processing. Finally, we analyze 3b and 1 separately, and we further separate neurons within each area by the relative position between their receptive field and the stimulation site (center [RF_1_], periphery of the center [RF_2_] and far away from the center [RF_3_]). Separating our data this way allows us to address a number of issues: a) the functional differences between these two areas, which are often overlooked, but which previous evidence has pointed towards (17), starting with the observation that each one has a complete somatotopic map of the body (18, 19); whether neurons present multisensory responses due to b) their receptive field, c) their area or d) the presence of functional clusters within each area. All these possibilities are obfuscated in studies with lower spatial resolution but have important implications: if primary somatosensory neurons respond to acoustic stimuli, for example, the foundational concept of receptive fields must be reconsidered.

In summary, our work analyzes the firing rate coding and variability to 1) uncover evidence of areas 3b and 1 carrying out multisensory processing, and 2) characterize the differences between responses in these areas. Our analyses were done at both the single neuron and population levels. Regarding the first point, the results that follow establish that these areas exhibit are uniquely unimodal responses to the principal modality and not the acoustic stimuli. Multisensory processing is a controversial subject for which some studies (3–13) have found positive results, while others (20, 21) yielded no evidence of such processing in primary sensory cortices. As for the second point, the results that follow establish that these areas exhibit distinct responses, with area 1 showing more variability, integration, and processing. Our study finds that, even though these areas differ in firing rate coding, latency, variability, and intrinsic timescale, both are only modulated by tactile stimuli. This leaves areas 3b and 1, components of S1, as unimodal.

## RESULTS

### Bimodal detection task

Two monkeys (*Macaca mulatta*) were trained to perform the BDT, which consisted of reporting the presence or absence of a vibrotactile (Tac) or acoustic (Ac) stimulus, presented in the range from sub- to supra-threshold intensities. Both types of stimuli had a fixed frequency of 20Hz and lasted 0.5s. For the tactile condition, amplitude of stimuli delivered to the skin was modulated while for the acoustic condition, the volume of a 1 kHz pure tone was adjusted. Both modalities were interleaved with an equal number of trials where no stimulus was delivered (Abs). Animals pressed one of three push buttons to report their choice: Tac, Ac, or Abs (Fig. 1A). Push buttons were located a reaching level in the left quadrant relative to the midline of the animal’s body. While monkeys performed the BDT, neuronal activity was recorded within in S1 (areas 3b and 1; Fig. 1C) and neurons were classified according to the relative position between stimulation zone and receptive field (Fig. 1D) (see Methods and Fig. 1D for RF definitions: RF_1_, RF_2_ and RF_3_). Psychometric measurements for both modalities showed that animals were trained to perform the task up to their psychophysiological thresholds (Fig. 1E).

**Figure 1.**
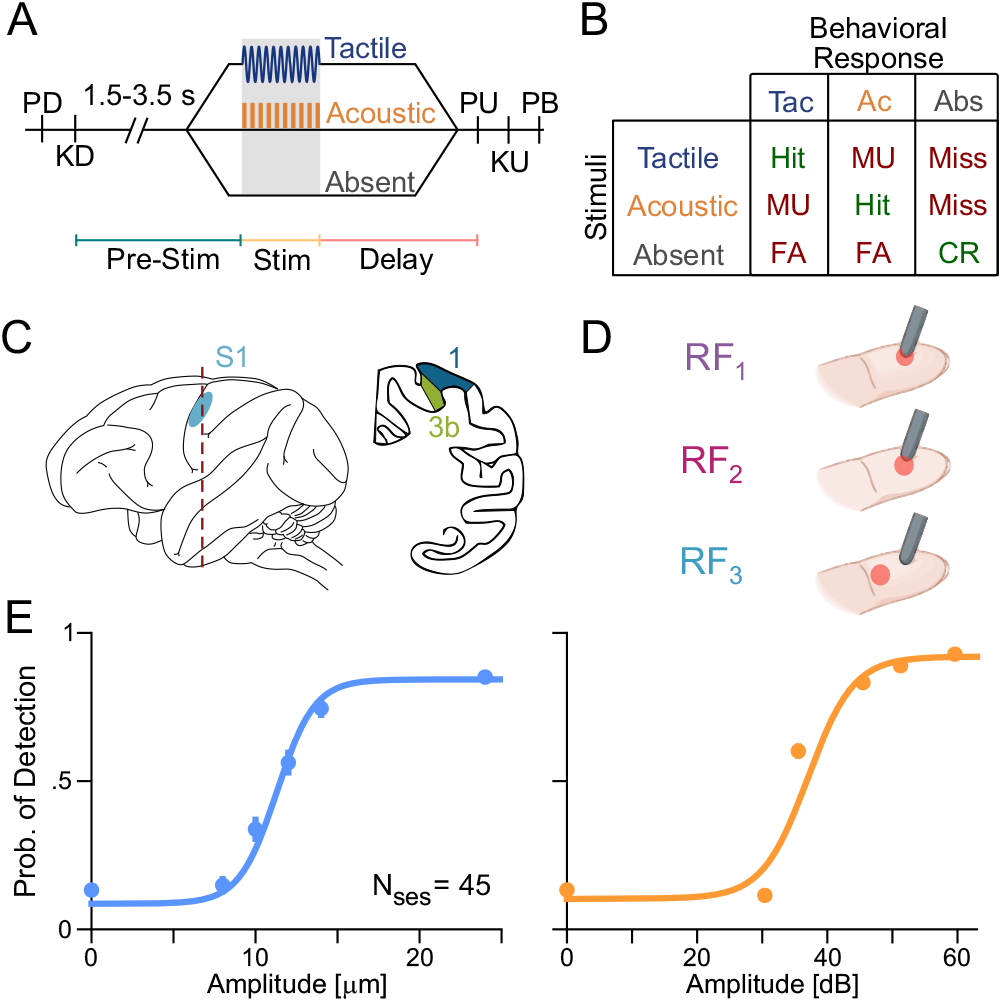
(A) Schematic representation for the bimodal detection task. Trials start when a mechanical probe indents the glabrous skin on one fingertip of the monkey’s restrained right hand (probe down event, “PD”). Immediately after PD, animals respond by placing their free left hand on an immovable key (key down event, “KD”). After KD, a variable period (1.5-3.5 s) is presented to prevent the animal’s prediction of stimulus onset. This is followed by a stimulus presentation (0.5 s), which can be tactile or acoustic (both 20 Hz), or absent. After stimulation, a fixed delay period (2 s) is presented followed by the probe up event (“PU”). This last event serves as the “go” cue for the monkey to release its hand from the key (key up event, “KU”) and report its decision using one of the three buttons placed in front of him, at eye level (push button event, “PB”). The three buttons indicate the following responses: “tactile stimulus present”, “auditory stimulus present” and “stimulus absent”. Correct responses were rewarded with a few drops of fruit juice. (B) Chart of trial outcomes according to stimuli versus behavioral responses. Correct responses are colored green and incorrect responses are colored red. In the absence of stimuli an incorrect “stimulus present” response indicates a false alarm (FA), while a “stimulus absent” response results in a correct rejection (CR). When a stimulus is present, a hit trial indicates a “present” response matching the given stimulus modality; a trial is misunderstood (MU) when the “present” response does not match the modality; finally, a trial is a miss if an “absent” response is given. (C) Illustration of the brain’s left hemisphere with S1 marked (left, light blue) and a coronal brain slice (right) with the areas 1 (blue) and 3b (green) highlighted. (D) Illustration of finger with receptive field locations (RF_1_, RF_2_, RF_3_) relative to the mechanical probe. (E) Psychometric curves for tactile (blue) and acoustic (orange) stimulation. Each modality had 5 classes of stimuli ranging from 0 to 24 μm or from 0 to 59 dB with a total of 6750 trials over the course of 45 sessions.

### Response properties of S1 neurons

We recorded activity from 67 neurons in area 3b and 313 in area 1, while monkeys performed the BDT. Based on recent evidence that suggests differences in their hierarchy (17), throughout this manuscript we analyzed neurons from areas 3b and 1 separately. For area 3b we found: 45 neurons (67.2%) that corresponded to RF_1_, 10 (14.9%) to RF_2_, and 12 (17.9%) to RF_3_. For area 1, 81 (25.9%) corresponded to RF_1_, 108 (34.5%) to RF_2_, and 124 (39.6%) to RF_3_. Fig. 2 and Fig. S1 show the responses of three individual neurons recorded in areas 1 (RF_1_ and RF_3_) and 3b (RF_2_) (and vice versa in Fig. S1, where an RF_2_ neuron from area 1 and RF_1_ and RF_3_ neurons for area 3b are displayed) with their corresponding normalized activity, during the tactile and acoustic trials. It is important to note that RF_1_ neurons faithfully represent intensity modulations of vibrotactile stimuli (Figs. 2A and S1A) while, despite belonging to the same brain area, this property is diluted in RF_2_ (Figs. 2B and S1B) and RF_3_ (Figs. 2C and S1C) neurons. Notice that RF_2_ neurons responded only during suprathreshold stimulation (dark pink). On the other hand, no activity modulation was found during acoustic stimulation regardless of the RF location or brain area, which suggests that the responses originating from cortical areas 3b and 1 are insensitive to acoustic stimuli, regardless of their stimulus intensity. To test this observation in the full population, we computed the normalized activity of all RF_1_ neurons during tactile and acoustic stimulation for both brain subareas (Fig. 3A and B). The results showed the same pattern as in the individual units, where neural responses in 3b and 1 were actively modulated in the tactile condition but not in the acoustic one. Similarly, as was observed in RF_2_ and RF_3_ single neurons, neural population activity diminished in both brain areas for these RFs (Fig. S2A and B) without showing any sort of modulation during acoustic stimulation were applied, no matter the intensity (Fig. S2C). Again, the RF_2_ population modulated its activity only with suprathreshold stimuli.

**Figure 2.**
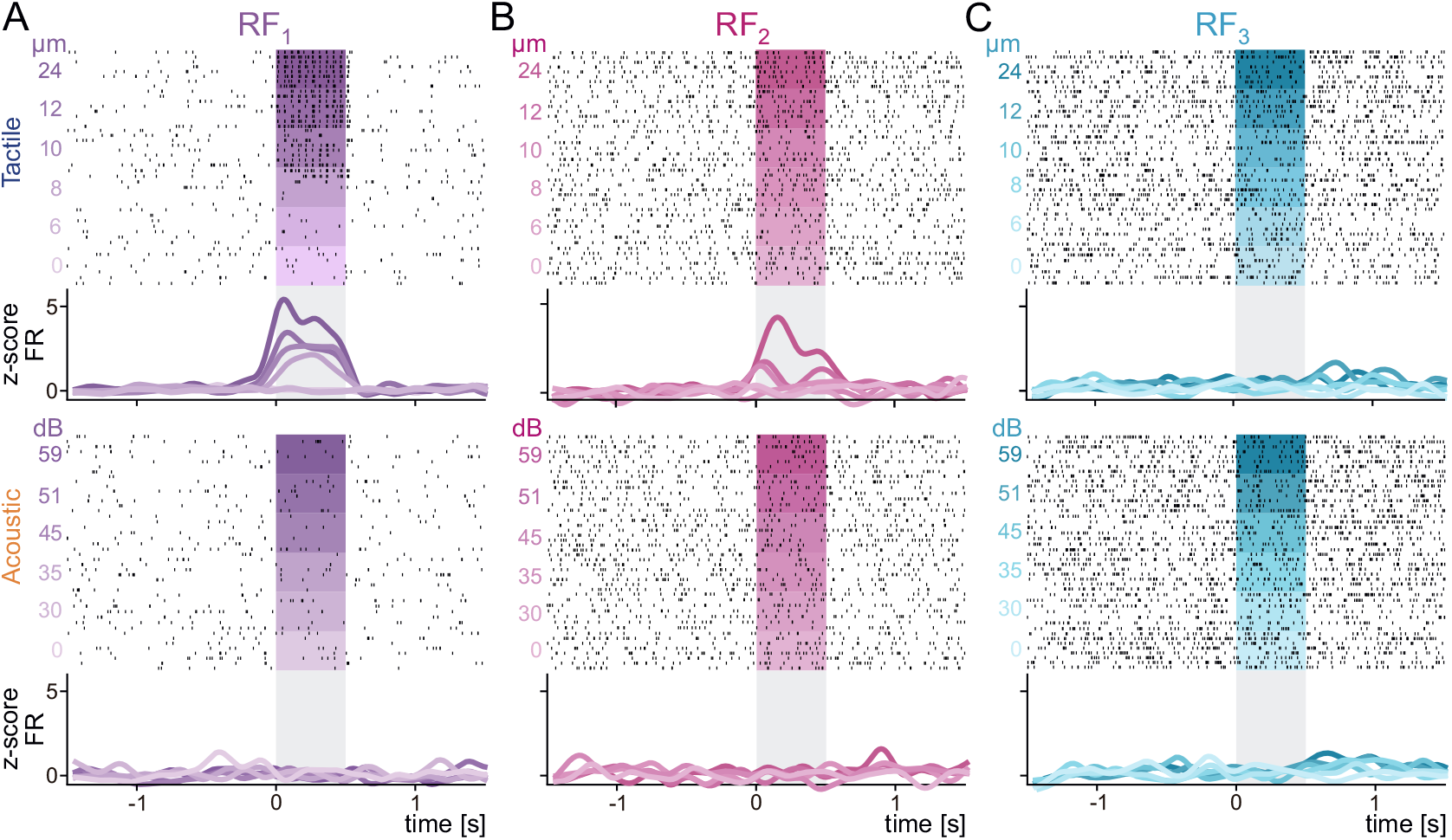
(A-C) Raster plots of three neurons recorded with different receptive fields. Normalized neuronal activity is shown below each raster. Neurons were recorded in area 1 (A and C panels) and 3b (B panel), for the three different receptive fields (RF_1_, purple; RF_2_ pink; RF_3_ blue), in the tactile (top) and acoustic (bottom) modalities. In the raster panel, black ticks represent neuronal spikes while colored rectangles represent the stimulation period. On the other hand, in the activity panel, colored lines represent the average of normalized neuronal activity obtained from trials with the same stimulus class and gray rectangles represent the stimulation period.

**Figure 3.**
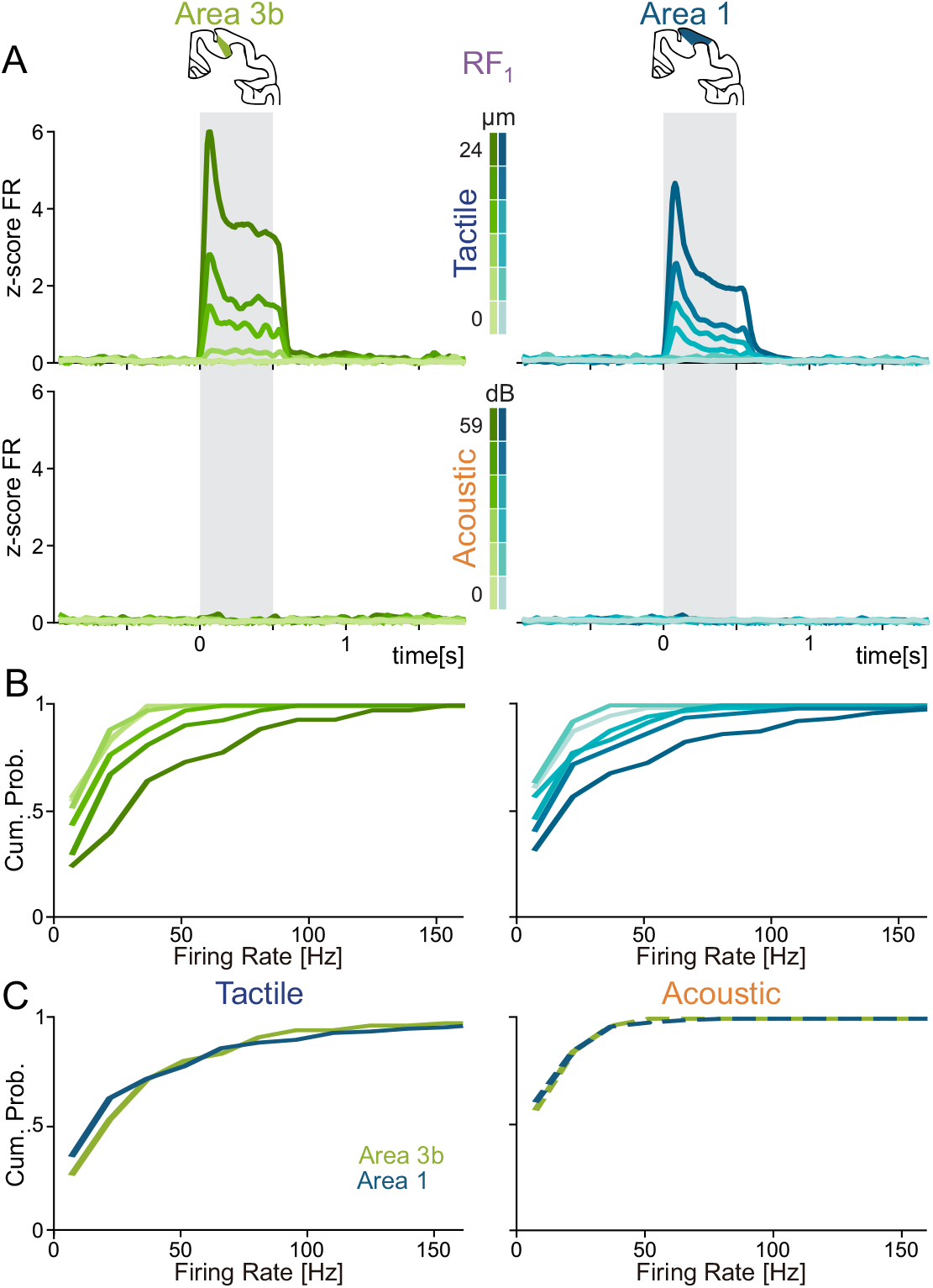
(A) Normalized population activity for the tactile (top) and acoustic (bottom) conditions, obtained from neurons recorded in areas 3b (left column) and 1 (right column) in the RF_1_ condition. Each colored line represents the average of normalized activity obtained from trials of the same class of stimulus. (B) Cumulative distributions of firing rate for the tactile modality calculated during the stimulus period for areas 3b (left) and 1 (right). (C) Cumulative distributions of the firing rate considering only the supraliminal tactile (left, 12 and 24 μm) and acoustic (right, 52 and 59 dB) stimuli, for neurons recorded in areas 3b and 1 (AUROC_tac_=0.545 ± 0.037, p=0.370; AUROC_acus_ = 0.516 ± 0.029, p=0.736).

These results suggest that neurons in areas 3b and 1 are unimodal. Nevertheless, we still wondered whether there was some slight modulation in the neural activity during the presence of acoustic stimulation. To further investigate such a possibility, we performed neurometric analysis with the neural activity data, using the optimum criterion technique. Given that sensory neurons are recognized for accurately codifying the frequency of stimuli (21, 22), we also computed the neurometric curves using the Fourier transform of the firing rate during the stimulation epoch (Methods). Figs. S3A and B show neurometric curves for neurons of areas 3b for the tactile and acoustic experimental conditions. In agreement with previous studies (16, 22), neurons from both areas yielded a strong and marked response when the tactile stimulation set was applied. Further reaffirming this behavioral response, periodicity neurometric curves showed similar results: no modulation for the acoustic stimuli, while in the tactile modality there was a detriment in the periodicity for the neurons from area 1 compared with those in area 3b. These results indicate that neurons from areas 3b and 1 respond to tactile stimuli while remaining unaffected by acoustic stimuli.

### Rate and periodicity as neural codes

To further test this hypothesis, we measured the information contained in the firing rate and periodicity of all RF_1_ neurons in areas 3b and 1 using Shannon’s mutual information (see Methods). We found significant differences in both rate and periodicity information when tactile stimulation was applied, while no such variations were found in the acoustic case (Fig. 4A). Reflecting its purely sensory nature, 3b neurons yield high values of periodicity information, while neurons from area 1 show only half the value for this metric. This loss of periodicity information suggests differences in processing between areas 3b and 1; being based on a rate rather than periodicity code. No significant differences (AUROC: 0.585 ± 0.049, p=0.1, n=5000 permutations) were found in the information that single neurons from areas 1 and 3b carried in their firing rate (Fig. 4C). In contrast, we found significant differences in periodicity (Fig. 4D) between both areas in the tactile modality (AUROC=0.708 ± 0.047, p=0.0002, n=5000 permutations). This reduction in periodicity representation between areas 3b and 1 has been previously found in a discrimination task (23), but not in a detection task.

**Figure 4.**
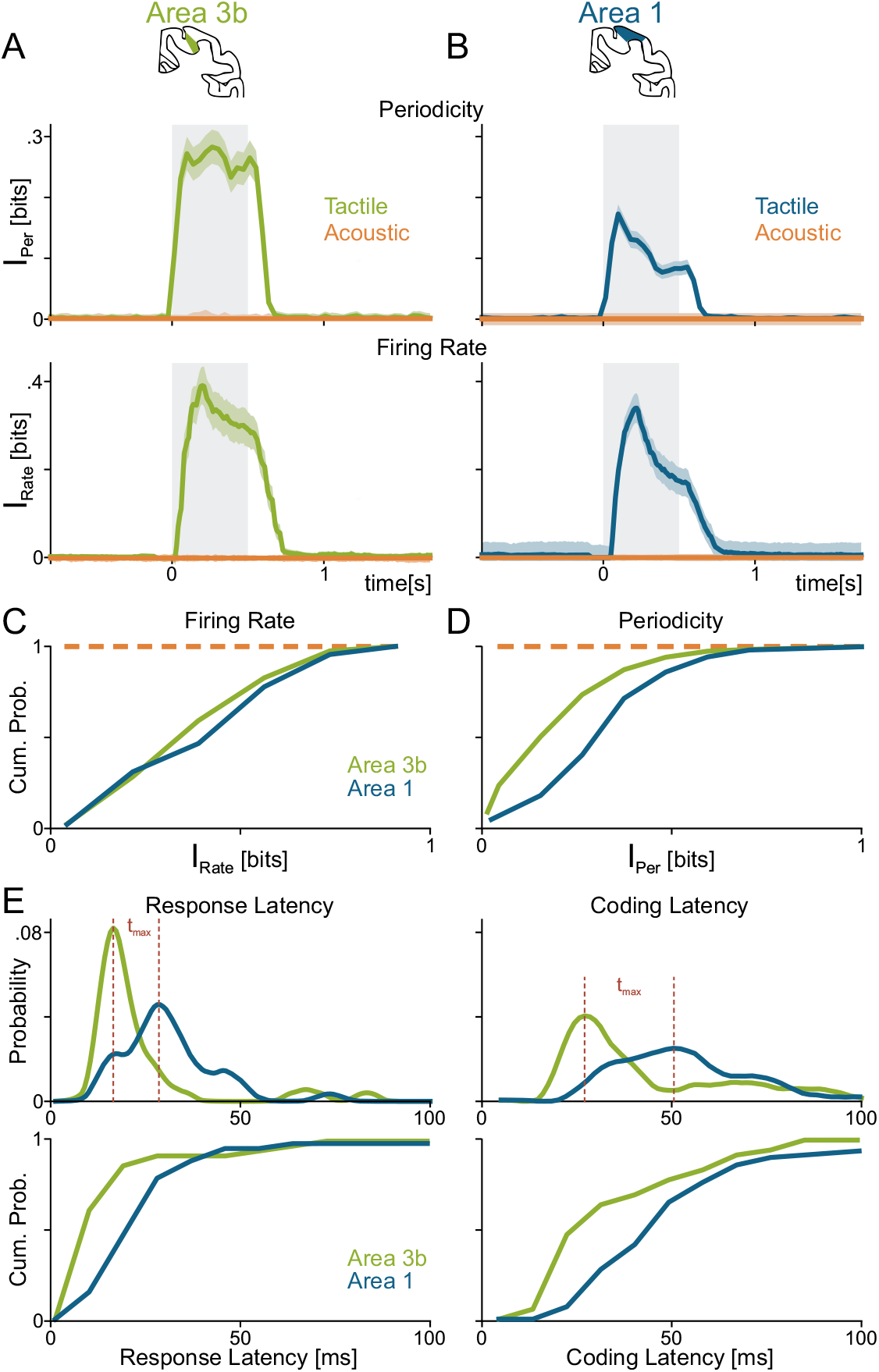
(A-B) Populational mutual information between stimulus intensity and (1) periodicity (top) and (2) firing rate (bottom). Both types of mutual information were calculated for tactile (areas 3b [green] and area 1 [blue]) and acoustic (orange) conditions for neurons recorded in areas 3b (A) and 1 (B). (C-D) Cumulative distribution of information, in the stimulation period and tactile modality, for rate (AUROC=0.59±0.049, p=0.1) and periodicity (AUROC=0.708 ± 0.047, p<0.0002). (E) Probability distributions (top) for response (left) and coding (right) latencies for the tactile modality in areas 3b and 1. The peaks of the probability distributions (t_max_) were 21.14 ms (area 3b) and 29.72 ms (area 1) for the response latency and 36.67 ms (area 3b) and 53.25 ms (area 1) for the coding latency. On the other hand, the bottom panel shows a comparison of the cumulative distributions between both areas for the response (AUROC=0.759 ± 0.050, p<0.0002) and coding (AUROC=0.711 ± 0.062, p=0.002) latencies.

For acoustic stimuli, no rate or periodicity information was found in either area (Figs. 4C and 4D, dashed orange line). This is consistent with the results shown in the previous section where acoustic inputs do not generate any sort of activity modulation and reinforces the idea of the unimodal nature of these primary sensory areas. Given the changes in periodicity information between areas 3b and 1 during tactile stimulation, we asked whether discrepancies were also present in their response latencies (RL) and coding latencies (CL). We observed that RL and CL distributions (Fig. 4E) were consistently left-shifted for 3b with respect to area 1. Quantification of tmax values for RL (21.14 ± 2.27ms, 3b; 29.72 ±1.76 ms, 1; upper left panel) and CL (36.67 ± 3.58ms, 3b; 53.25 ±3.38 ms, 1; upper right panel) confirms this observation, exhibiting significantly smaller tmax values for 3b with respect to 1. We verified the significance of this shift left by computing the AUROC values across areas 3b and 1 distributions for RL (0.759 ± 0.05, p<0.0002; lower left panel) and CL (0.711 ± 0.06, p=0.0002; lower right panel). These results strongly suggest that neurons from 3b are faster to respond and code the identity of tactile stimuli than those in area 1. Although differences in intrinsic timescales were recently identified (17), as far as we know this is the first evidence that shows a marked difference in the processing of vibrotactile stimuli between these two sensory areas.

### Variability fluctuations during the tactile and acoustic stimuli

Considering that neither firing rate nor periodicity have shown changes during acoustic stimulation, we wondered whether neural variability was modulated by acoustic stimuli. The population’s distribution of Fano Factor (FF) values showed no changes in either subarea during acoustic stimulation (Fig 5A and 5B). However, when tactile stimulation was presented, a clear decrease in variability (greater in area 1) was observed in both areas, highlighting once again the unimodal nature of these sensory areas. While this decrease was previously reported in visual areas (24), to our knowledge this has not been shown to occur in areas 3b and 1. Further, these results are noteworthy given that previous studies have suggested that variability modulation could be an indicator of the presence of a stimulus of another modality (25).

**Figure 5.**
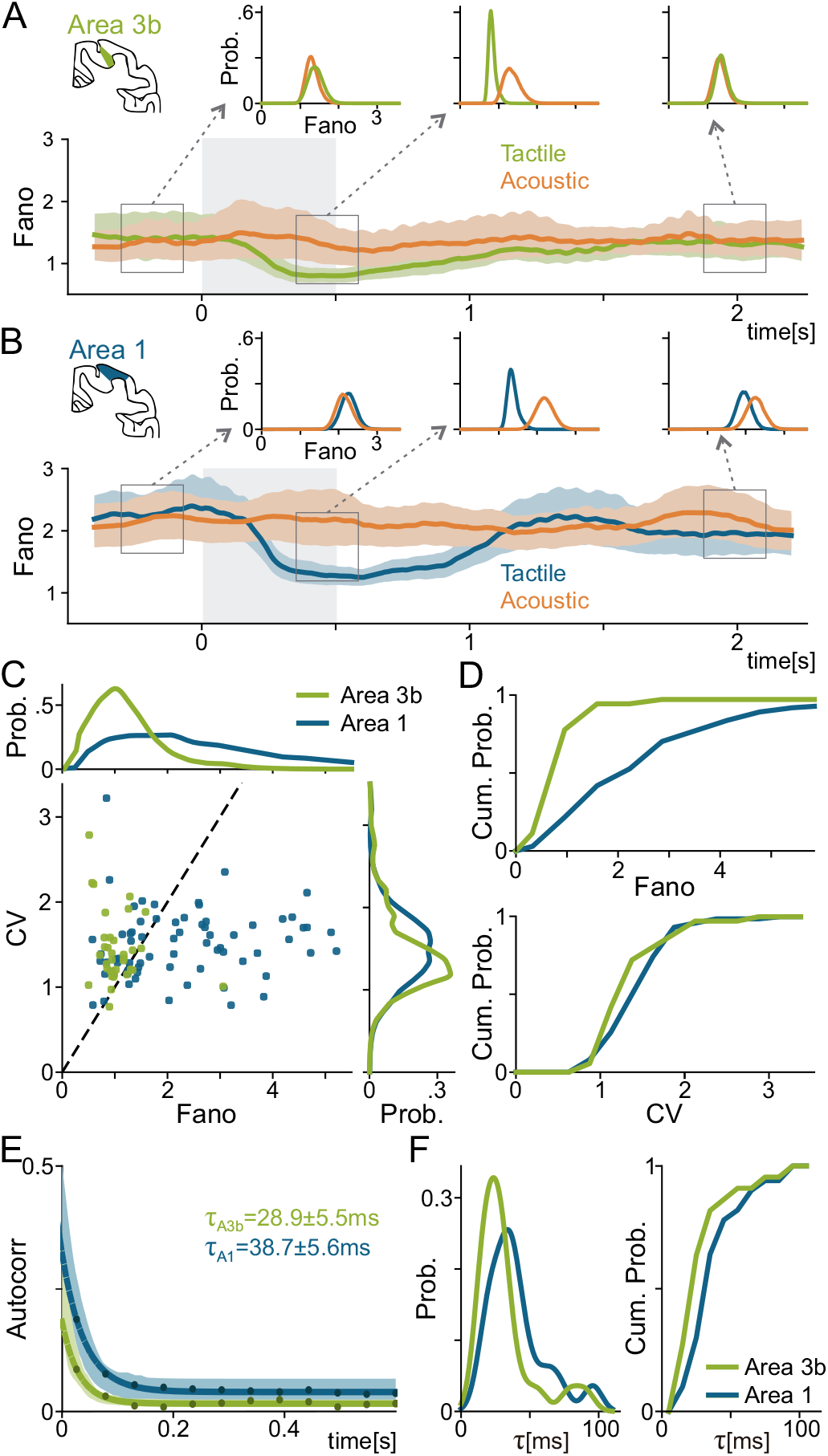
Populational Fano factor (FF) in time dependent manner for area 3b (A) and 1 (B). FF was calculated for both modalities: tactile (green [area 3b] and blue [area 1], traces) and acoustic (orange traces). In both areas and for every modality, a set of probability distributions were calculated during a short time interval indicated by empty gray squares. Intervals displayed include the pre-stimulus period (left), the end of the stimulus period (center) and the delay period (right). (C) Scatter plot comparing the coefficient of variation (CV) vs FF, during the pre-stimulus period for areas 3b and 1, accompanied with its corresponding probability distributions. (D) Comparison of cumulative distributions obtained for areas 3b and 1 of FF (top, AUROC=0.758 ± 0.035, p<0.0002) and CV (bottom, AUROC=0.538 ± 0.038, p=0.478). (E) Populational autocorrelation functions yielded timescales of 28.9 ± 5.5 ms for area 3b and 38.7 ± 5.6 ms for area 1. (F) Probability (left) and cumulative (right) distributions of the timescale constants obtained in the pre-stimulus period for area 3b (green) and 1 (blue).

The variability differences between areas 3b and 1, during the stimulus period, raised the question of whether these two areas are intrinsically different at the fluctuation level. To explore such a possibility, we measured the FF and the coefficient of variation (CV) during the pre-stimulus period. Noticeable differences between area distributions were detected, with significantly higher intrinsic variability in area 1 with respect with area 3b (AUROC=0.758 ± 0.035, p<0.002, n=5000 permutations) (Fig. 5C & 5D). These results gave us the idea of the existence of a clear separation in how areas 3b and 1 process the identity of the tactile stimuli, suggesting a hierarchy in the circuit of tactile information processing, where area 3b will be then followed by area 1. Moreover, this suggests that neurons from area 1 integrate responses from more neurons than area 3b, giving rise to larger receptive fields (18) and higher variability.

To further confirm this hypothesis, we computed the firing rate autocorrelation analysis by using the neurons in both areas (Fig. 5E) (26, 27). Remarkably, we observed that the decay time constant (τ) was smaller for area 3b than area 1. This can be interpreted as area 1 has a greater amount of reverberations at the network level, i.e. a major level of integration (28). In summary, the results displayed up to this section provide direct evidence that areas 3b and 1 unimodally process stimuli information. Furthermore, they reveal a processing hierarchy between these two areas, with tactile stimuli information being integrated differently in area 3b than in area 1.

### Are there activity clusters differentiating areas 3b from 1?

At this point, we wondered whether the differences between the responses of neurons in areas 3b and 1 were significant enough to distinguish to which area they belong to. In other words, would single neurons create two non- or low-overlapping clusters representing the two areas? Or is there an overlapping continuum of responses that connects both areas? To address this question, we performed a non-linear dimensionality reduction on the concatenated activity profiles. The UMAP (Uniform Manifold Approximation and Projection) algorithm considers similarities at both the local and global scales (29). For our data, the first UMAP dimension shows an apparent separation between the activity of both areas (inset distribution Fig. 6A). However, to test this observation, we computed a density based metric to find clusters (30). With this procedure, we determined the existence of two density peaks (Fig. S4A), one corresponding mainly to area 1 and the other to area 3b. Therefore, our analysis suggests the existence of two clusters of neurons determined purely by their responses to a set of stimuli. Based on this initial analysis, we fit a non-linear classifier with support-vector machine (31). Fig. 6B shows the mean and the standard deviation of the activity profiles by using the best iteration of the classifier (0.79 ± 0.27% mean of cross validation by using 35% of the dataset as testing). A high classification accuracy is obtained despite the clusters’ unclear visual appearance, suggesting that the responses are gradually transformed from area 3b to 1. Pronounced peaks at the beginning of the stimulus period are dampened and responses to sub-threshold stimuli gradually increase when transitioning from area 3b to 1 (Fig S4 B). This transformation may be related to the differences in timescale integration observed in Fig. 5. This mirrors the relationship between S1 and the ventral posterior lateral nucleus of the thalamus (VPL)(32), with area 1 neurons appearing more stable (in their coding) than those in area 3b. Even if our results are not based on and do not imply an anatomical hierarchy, our findings could lead to a more refined hierarchy of touch processing: VPL -> 3b -> 1.

**Figure 6.**
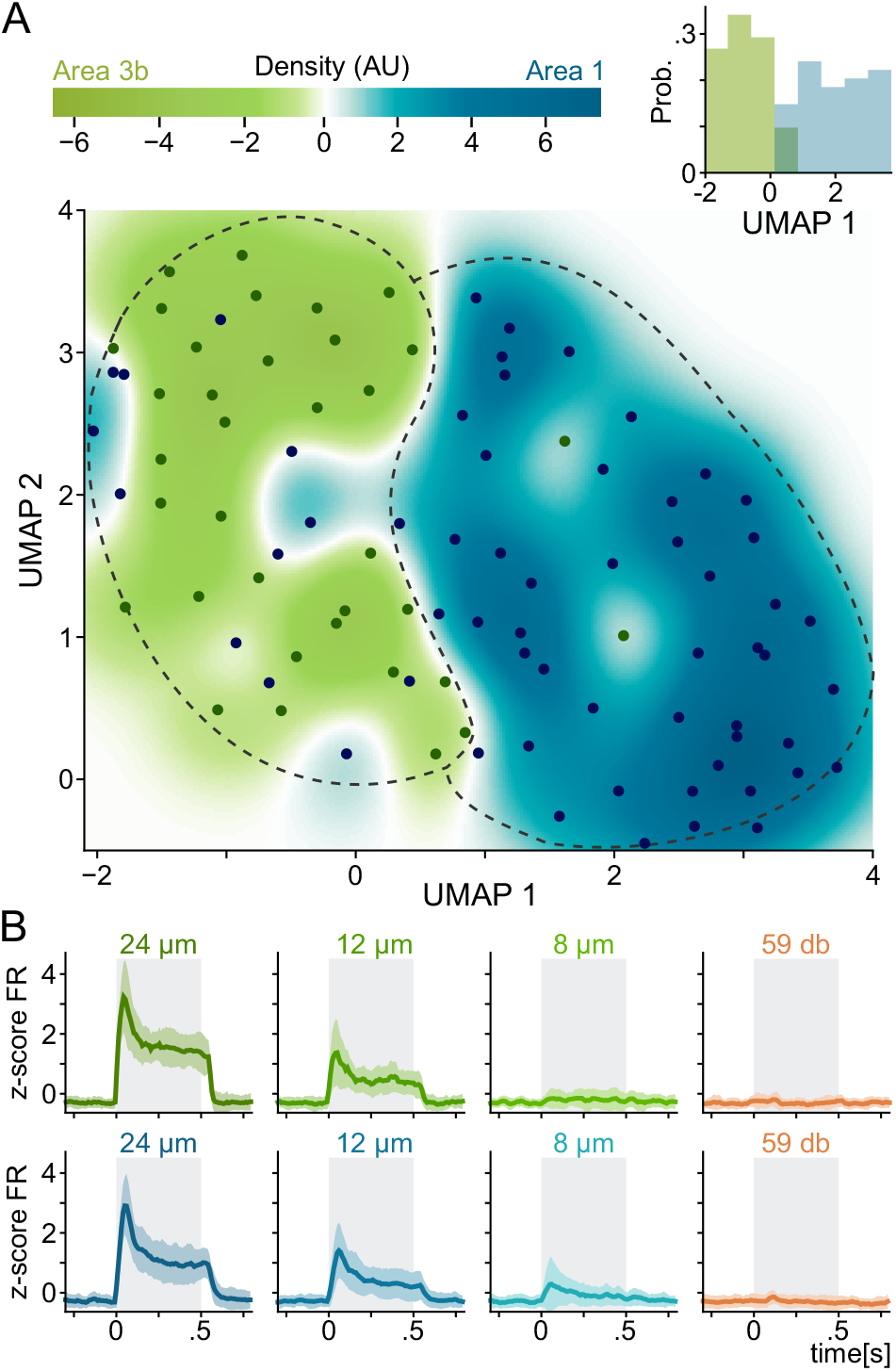
(A) Density of 2-D non-linear projection of the normalized neural activity obtained with UMAP (29). Green regions exhibit dominance of neurons from area 3b while blue regions show dominance for area 1. Histogram inset shows the probability distribution of each area in the first UMAP dimension. (B) Average normalized firing rates of neurons classified with a non-linear SVM (31). Note that most 3b neurons reside in class 1 (top) and area 1 neurons in class 2 (bottom). Firing rate profiles include the supraliminal (24 μm), liminal (12 μm) and subliminal (8 μm) ranges. Acoustic profiles for the supraliminal condition (59 dB), in both areas, are shown in orange. While class 1 modulated its activity with 8 μm, class 2 did not.

## DISCUSSION

To investigate the existence of multisensory processing in S1, we recorded the neural activity of subareas 3b and 1 during a detection task where hearing and touch compete. By separating single units according to the relative position between their receptive field and the stimulation zone (RF_1_, RF_2_ and RF_3_), we could study the response and variability of neurons whose receptive field was not directly stimulated. At the unit and population levels, we found that both areas process and encode the tactile but not the acoustic stimuli. Yet, we found three significant differences between areas 1 and 3b: 1) a decrease in periodicity information from area 3b to 1, suggesting that neurons from area 1 have a less faithful representation of the stimuli; 2) an increase in variability and timescale from area 3b to 1, indicating differences in their information processing; 3) neural responses suffer a gradual transformation between areas 3b and 1. Particularly, we observed that neurons from area 1 responds stronger to subliminal stimuli than neurons from area 3b, highlighting the differences in processing between the areas. Our results suggests that neurons from areas 3b and 1 are unimodal in their sensory processing and reveals the existence of a novel intrinsic level of processing-based hierarchy within S1.

The main characteristic of primary sensory cortices is their representation of features from the external world. Thus, to consider a response as overtly multisensory it should code the properties of stimuli from more than one modality. In contrast, our work reveals that the rate and periodicity of areas 3b and 1, at the single and populational levels, encode information unimodally. Furthermore, we found that periodicity information decreases from 3b to 1, even though these areas showed similar firing rate responses. Previous studies in a different task have shown that, while neurons from area 3b display phaselocking and periodic responses (33), the secondary somatosensory cortex (S2) also demonstrates a loss of periodicity information at the population level (23) with the construction of more complex and abstract coding (34). The role of S2 in the BDT is an important avenue for future research. Moreover, as stated in the Results, variability analysis is a key concept that should be considered and has rarely been applied in the context of multisensory processing.

Even if isolated neurons demonstrate stochastic variability they are much more reliable than those observed in brain recordings (35). Ergo, at least part of the cortical variability comes from fluctuations in synaptic inputs, shaped by the network’s features (36, 37). In the mammalian brain, the degree of variability at the single-neuron level increases with the stages of sensory processing, being lowest in the periphery and highest in cortical structures (27, 38). Understanding the nature and origin of variability is imperative to the study of the neural codes used for the representation and processing of information in cortical networks (39–41). In relation to our subject of interest, a previous study has proposed that multisensory activity could be manifest in firing rate fluctuations (25). Nevertheless, we found no significant Fano Factor modulation during acoustic stimulation. Remarkably, the activity of areas 3b and 1 showed consistently significant differences between their variabilities during tactile stimulation. Additionally, both areas also differed in their responses and coding timescales. Taken together, these results support the existence of a hierarchy in the processing of information between areas 3b and 1.

At this point, we asked ourselves whether these two populations are different enough to reliably identify to which one a neuron belongs. To answer this, we leveraged the topological properties of the firing rates of neuronal populations using a novel manifold learning method (UMAP) (29). In other reports, this algorithm has revealed interesting dynamic and geometric properties (42, 43). Our analysis revealed a clear separation of areas 3b and 1, which is nonetheless connected by a continuous transition between their neural activity profiles. This transition ignored the acoustic stimulus completely, considering only the tactile responses, particularly to low amplitude stimuli. The discrepancy in timescales across these two subareas may be one of the network features that promote this separation. Area 1 neurons are equipped with larger receptive fields and timescales compared to neurons from area 3b, which may cause greater sensitivity to low amplitudes via sensory integration. Future simultaneous recording studies across this network should measure information flow (44) to look for more evidence to solve this question.

Historically, sensory processing in area 1 has been considered indistinguishable from that in area 3b. In our opinion, this conception is due to their similar firing rate patterns and a tendency to focus on single units rather than populations. The results shown here provide strong evidence that areas 3b and 1 are located at different levels in the somatosensory network’s processing hierarchy. This is in line with recent results, where analogous distinct steps in the processing and abstraction of visual information were found (27). However, disparities in thalamocortical input between the areas provide an alternative explanation that is not accounted for in our work. Future research should focus on answering whether the processing differences found here emerge as a consequence of their network features or due to discrepancies in the amount of information projected from the thalamus.

Contemporary research in which different types of brain signals were used, such as LFP (8, 9), EEG (5, 13) or fMRI (12, 45), has tried to demonstrate that multisensory activity appears at the early stages of processing in the primary sensory cortices. Up to this point, the reader can see that the results presented in this work differ notoriously from those; we think that this difference could be due to multiple factors.

First, our analysis does not explore the aggregated activity of neurons as LFP or EEG do. So, if the aggregated or synaptic activity of S1 encodes some properties of acoustic stimuli might still be debated. Multisensory processing could be present in facets of the network outside the scope of our data, which could be detected using different approaches: 1) studying correlated variability at the level of spikes and/or oscillations; 2) taking LFP approaches which are common in the study of the attention phenomena and the transition between network states (46); And finally 3), if multisensory processing is present in the interaction between areas, this could be measured through phase or amplitude coherence, joint fluctuations or interaction metrics with any combination of spikes and fields.

Second, our task is competitive in nature; it divides attention between two senses, with their presence being mutually exclusive. This contrasts with most cognitive tasks employed in other studies, where the stimuli are synchronized and cooperative. If multisensory activity only appears in early sensory cortices during cooperative stimulation, then it might be more appropriate to conceptualize it as an effect enhancing unimodal processing. Additionally, it is important to consider that the use of two synchronized stimuli hinders neural coding analyses due to the introduction of more variables into the coding problem. This multivariate extension is sometimes not well controlled or does not take the coding problem into account when designed, possibly yielding uninterpretable results (47).

Finally, on the subject of coding, experiments described in other works lack control at the finest level of the stimuli’s physical parameters (5, 8, 9, 12, 13, 45). This makes it impossible to determine which physical properties were being encoded by neurons, which would have been indisputable proof of sensory modulation. In contrast, our experimental framework was designed to look for the most specific form of multisensory processing possible, with the physical properties of stimuli accurately controlled. In addition, performance in the BDT is scarcely impacted by dividing attention (15), making it possible to isolate the response to pure unimodal stimuli and determine unequivocally the effects of another modality in S1.

The opening question of the Introduction section, about a car’s radio, is most often used as an example of attention. As mentioned briefly there, attention gives an alternative hypothesis to multisensory processing; attentional mechanisms could contribute to the differences between neural responses in unimodal and those in multimodal tasks. We also mentioned that our task should not be affected by these mechanisms (15). Still, it is relevant to discuss some of the findings that relate attention to the topics of our study. For example, attentional effects have been evaluated in circuits near the primary sensory areas, but it remains unclear how attentional phenomena affect primary processing (48, 49). It has also been reported that multisensory processing could be task-dependent (50, 51). This could point to more elaborate interactions between the hypothetical early multisensory integration, attention, and task demands. The study of these interactions could greatly benefit from the synthetic approach advocated in (52); there, the concept of attention was critiqued for being too broad for the study of neural systems. Rather, the approach advocated involves proposing specific stimuli selection processes relevant to behavior. In any case, the experiment presented here is not optimal for the study of attentional effects. For this purpose, a better experimental design based on the BDT could consist of adding two more stimuli: one serving as a cue, prior to the original stimulus, indicating to the subjects which modality to attend; and the other, synchronized with the original, acting as a distractor. With this task, it would be possible to disentangle the effects of attention, uncertainty, and multisensory processing in the activity of the primary sensory cortices, while also increasing the competition between modalities.

In brief, we did not find evidence in favor of acoustic processing at the level of firing rate nor variability for areas 3b and 1 of S1, no matter the relative location of the neuron’s receptive field stimulated during the BDT. Remarkably, we found clear differences between areas 3b and 1 at the level of variability, periodicity information and timescales. In addition, utilizing a powerful nonlinear dimensionality reduction technique we found a clear, but continuously connected, separation between these two areas.

## ACKNOWLEDGMENTS

We thank Gabriel Diaz-deLeon for his technical assistance. This work was supported by grants PAPIIT-IN205022 from the Dirección de Asuntos del Personal Académico de la Universidad Nacional Autónoma de México (to R.R.-P.) and CONACYT-319347 (to R.R.-P.) and CB2014-20140892 (to R.R.) from Consejo Nacional de Ciencia y Tecnología. S.Pa. (fellowship CONACYT-631990), H.D. and J.Z. are doctoral students from Programa de Doctorado en Ciencias Biomédicas, UNAM. L.B. is a postdoctoral student (Postdoctoral fellowship CONACYT-838783).

## MATERIALS AND METHODS

### Bimodal Detection Task

In this cognitive task, two monkeys (*Macaca mulatta*) were trained to detect the presence or absence of a single vibrotactile or acoustic stimulus (Fig. 1A). Stimulus intensities were varied from 6-24 μm and 34-59 dB for the tactile and acoustic modalities respectively. These ranges include sub-liminal, liminal and supra-liminal stimuli for both modalities. Vibrotactile stimuli were delivered to the skin of the distal segment of one digit of the monkey’s restrained hand, via a computer-controlled stimulator (BME Systems; 2-mm round tip) while acoustic stimuli were delivered by a speaker. Both stimulus modalities consisted of a sinusoidal at a frequency of 20 Hz. In the acoustic modality, each stimulus signal was composed of a pure tone (1 kHz), while in the vibrotactile modality each stimulus consisted of a train of mechanical-sinusoidal pulses. Both stimulus modalities were interleaved with an equal number of trials where no stimulus was delivered. Animals pressed one of three buttons to indicate their response: vibrotactile-stimulus-present, acoustic-stimulus-present, or stimulus-absent. Monkeys were rewarded with a drop of liquid for correct responses. Performance was quantified through psychometric techniques (Fig. 1E). Both animals were handled according to institutional standards of the US National Institutes of Health and the Society for Neuroscience. All protocols were approved by the Institutional Animal Care and Use Committee of the Instituto de Fisiología Celular of the National Autonomous University of Mexico (UNAM).

### Recording sessions and sites

Neuronal recordings were obtained with an array of seven independent, movable microelectrodes (1.5–3 [MO]) inserted into S1 (53). Recordings in both monkeys were made in areas 3b and 1 (Fig. 1C), contralateral to the stimulated hand. The location of electrodes was confirmed using standard histological techniques. Neurons’ receptive field locations were determined during the experiment by carefully stimulating different regions in the right restrained hand and observing its response. Neurons were classified into three categories according to their receptive field (RF) location: RF_1_ is composed of neurons whose RF was inside the stimulation zone (vibrotactile probe). Such neurons were characterized by showing a strong response when the probe touched the glabrous skin of one finger digit; RF_2_ is composed of neurons whose RF was close to the stimulation zone and were characterized by exhibiting a mild response when tactile stimuli were presented. Finally, RF_3_ is composed of neurons whose RF was located far away from the stimulation zone. Such neurons were identified by eliciting a null response, remaining unaltered, during the tactile stimulation.

### Data analysis

#### Firing Rate

For each neuron, we calculated a time-dependent firing rate for each trial. The firing rate was calculated by using rectangular causal windows with a standard window length of 50 ms and step of 10 ms. A normalized version (zscore) was calculated by subtracting the mean and dividing by the standard deviation of the firing rate estimated in the 1.5s interval previous to stimulus onset (33).

#### Neurometric curves

To calculate response distributions and the neurometric detection curves of S1 neurons, from each trial we obtained the firing rate during the stimulation period. We generated two groups from the recorded data, the first considering absent and tactile trials and the other where absent and acoustic trials were considered. For each group, neurometric curves were calculated as the proportion of trials in which the maximum firing rate reached or surpassed a criterion level (16). For each neuron, this criterion was chosen to maximize correct trials (the number of hits and correct rejections). From logistic fits, we calculated psychometric and neurometric detection thresholds as the probability that the proportion of ‘yes’ responses would be 0.5. The periodicity neurometric curve was calculated by defining a periodicity coefficient obtained during the stimulation period. Such a metric was calculated by estimating the firing rate using a causal kernel with a resolution of 100 Hz. Afterwards, we subtracted the mean and used a Hamming window. Then, we extracted the power of the frequency window from 18 to 22 Hz to obtain the mentionated coefficient. This periodicity coefficient was used to construct a neurometric curve by using the optimum criterion technique described above (21).

Mutual Information. For each neuron, we calculated two fundamentally different types of mutual information in a time-dependent manner: firing rate information (*I_RATE_*) and periodicity information (*I_PERIOD_*). *I_RATE_* was calculated with a firing rate (*r*) obtained with a causal window length of 200 ms and a step of 50ms. Then, we extracted the firing-rate distributions by stimuli group (*P(r|s)*) and the global probability distribution (*P(r)*). All distributions were estimated by time-bin. Then *I_RATE_* mutual information was estimated by using Shannon’s equation in a time dependent manner (23).

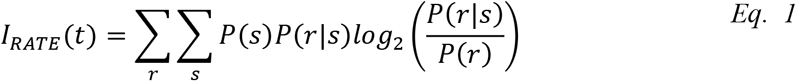

On the other hand, periodicity information was calculated in the following manner: we calculated a periodicity coefficient as a function of time using a mobile window of 200 ms and short-time-Fourier Transform (23). To compute the coefficient, we employed the firing rate using a causal kernel with a resolution of 100 Hz. Afterwards, we subtracted the mean and used a Hamming window. Then, we extracted the relative power of the frequency window around 20 Hz to obtain such periodicity coefficient (*perc*). This procedure resulted in a set of periodicity coefficient (*perc*) distributions for each stimulus intensity as a function of time (*P(per_coef_|s)(t)*). With this data, we calculated *I_PERIOD_* between periodicity at 20Hz and stimulus intensity by using Shannon’s equation:

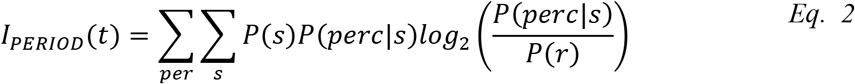

Information estimates were corrected by a standard shuffling procedure (54). Significant time-bins of information were determined by a permutation test (p=0.01, 1000 iterations) and corrected for multiple comparisons with a clustering method (55). Additionally, a populational information analysis for *I_RATE_* and *I_PERIOD_* was performed in a similar fashion by using the normalized firing rate (34). These estimations were corrected by the shuffling procedure followed by the asymptotic Panzeri-Treves correction (56).

#### Fano Factor (FF)

We employed a standard procedure to calculate the FF (24). For each neuron, we calculated the mean and variance of the normalized firing rate per time-bin for each stimulus intensity and modality. Afterwards, we pooled the mean and variances for each time-bin of all neurons grouped by brain area (3b or 1) and RF_1_. The Fano factor was estimated as the slope of the best linear fit of such responses for each modality.

#### Coefficient of Variation

To estimate the variability in the inter-spike interval (ISI) for each neuron, we separated neurons according to their brain area (3b or 1). Then, we selected the activity at the prestimulus period and calculated the distribution of ISI values. With those distributions, we then calculated the coefficient of variation as in the following expression:

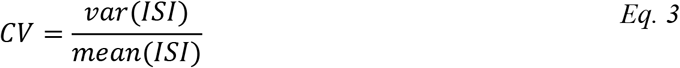

#### Autocorrelation

To characterize the autocorrelation of spike counts, we pooled the activity of neurons in the prestimulus period and grouped them by brain area (3b and 1). We followed the same methodological procedure as in (26) for single neurons. The basal period (−1 to 0 sec) was divided into successive time bins of 40ms duration and 20ms step. Then, for two-time bins separated by a time lag *t*, we calculated the across-trial correlation between spike counts *N*. Afterwards, this autocorrelation function of the time lag *t* between bins was fit (least-square procedure) by an exponential decay with an offset:

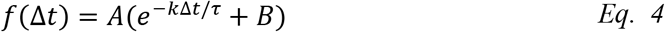

Where *τ* (tau) represents the decay rate constant, which measures the intrinsic population timescale. The offset (B) represents the contribution of timescales much longer than our observation window (26). We fit Eq. 4 to the full autocorrelation data from all neurons and trials. Hence, fits were performed at the population level and single-neuron level. Additionally, the estimation of the error of fit parameter *τ* (tau) was performed by population bootstrap analysis. Synthetic datasets were created by a resampling process of the neural population. Each bootstrap sample population is taken from the original by using sampling with replacement; each neuron may be repeated or absent in each bootstrap sample. Afterwards, a new fit was calculated for every new dataset, giving rise to a distribution. The 95% confidence interval of the distribution was computed to estimate the error in *τ* (tau) estimation.

#### Activity classification

We classified neural activity by concatenating the average of the normalized firing rate for a subset of stimuli amplitudes from the tactile (24 μm, 12 μm, 8 μm) and acoustic (59 db) modalities. Then we used Uniform Manifold Approximation and Projection (UMAP)(29), a novel manifold learning technique for dimensionality reduction, to find a 2D representation of the data that preserves its local and global topological properties. To corroborate the existence of isolated groups, we performed density based clustering analysis (30). In brief, we characterized each point by its local density by using a Gaussian kernel and its distance to the nearest point with a greater value. Next, by sorting such a metric, it is possible to determine the number of groups or clusters in the dataset. Based on the number of groups detected, we classified the data by using a non-linear support vector machine (SVM) (31). To evaluate the classifier performance, we executed a cross-validation test by selecting 35% of the data as a test sample set.

---

**Figure S1.**
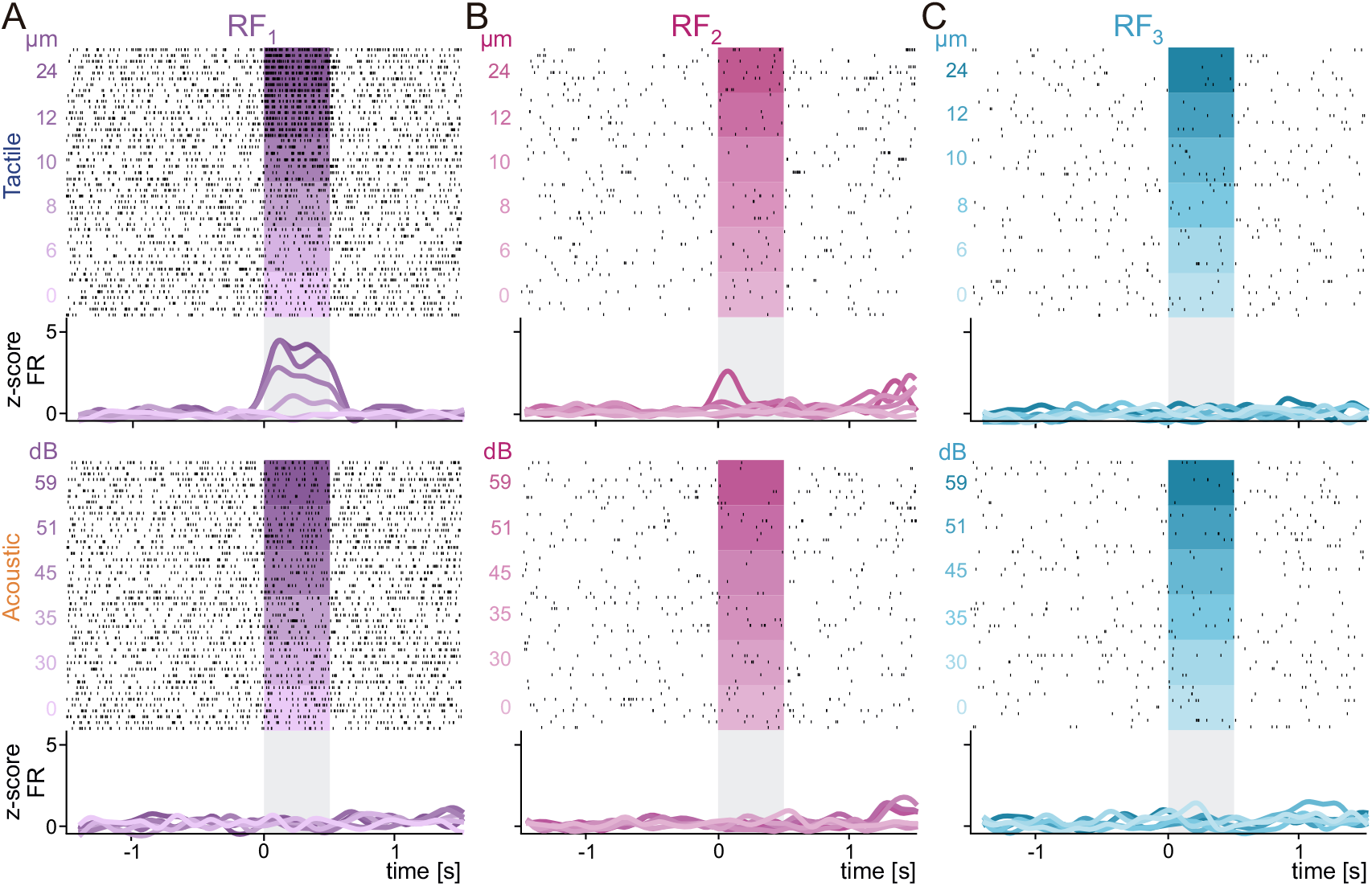
(A-C) Raster plots of three neurons recorded with different receptive fields. Normalized neuronal activity is shown below each raster. Neurons were recorded in area 3b (A and C panels) and 1 (B panel), for the three different receptive fields (RF_1_, purple; RF_2_ pink; RF_3_ blue), in the tactile (top) and acoustic (bottom) modalities. In the raster panel, black ticks represent neuronal spikes while colored rectangles represent the stimulation period. In the activity panel, colored lines represent the average of normalized neuronal activity obtained from trials with the same stimulus class and gray rectangles represent the stimulation period.

**Figure S2.**
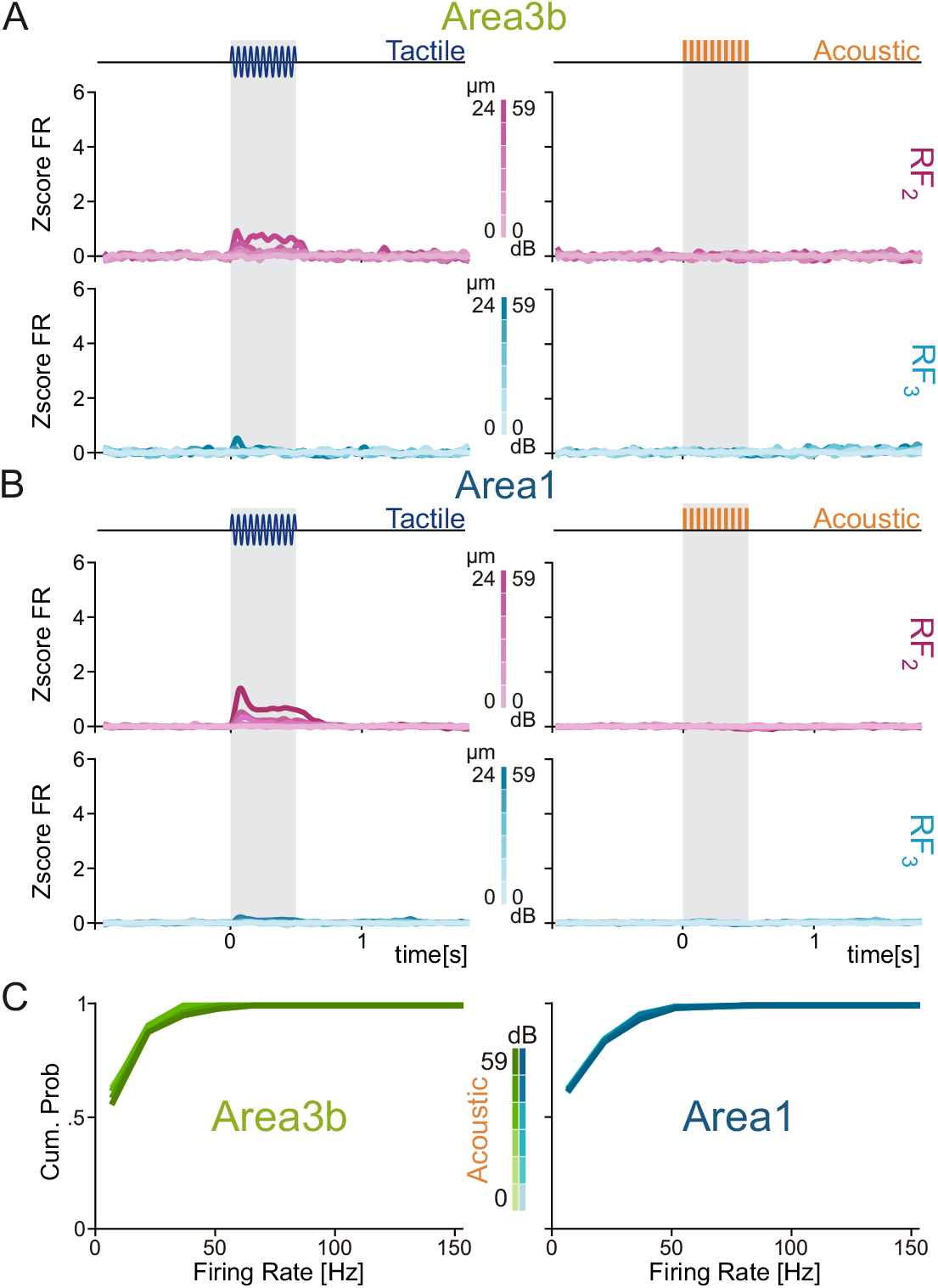
(A-B) Normalized neuronal activity (colored traces) in the tactile (left) and acoustic (right) modalities for the RF_2_ (pink) and RF_3_ (blue) of recorded neurons belonging to areas 3b and 1. (C) Firing rate cumulative distributions calculated for neurons from areas 3b (left) and 1 (right) during the stimulus period in the acoustic modality for the RF_1_ condition.

**Figure S3.**
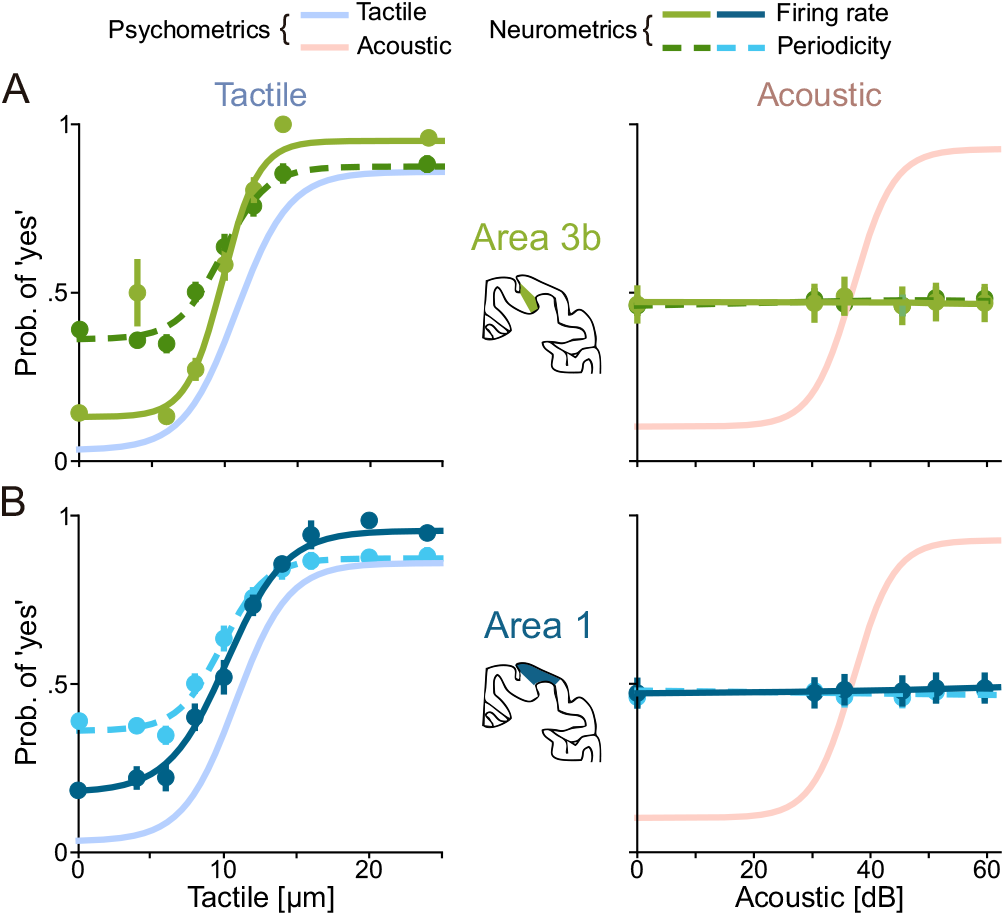
(A-B) Neurometric curves for areas 3b (green) and 1 (blue) in the tactile (left) and acoustic (right) modalities were calculated by using the firing rate (solid line) and periodicity (dashed line). Psychometric curves for both modalities are shown in a semi-transparent fashion on each panel.

**Figure S4.**
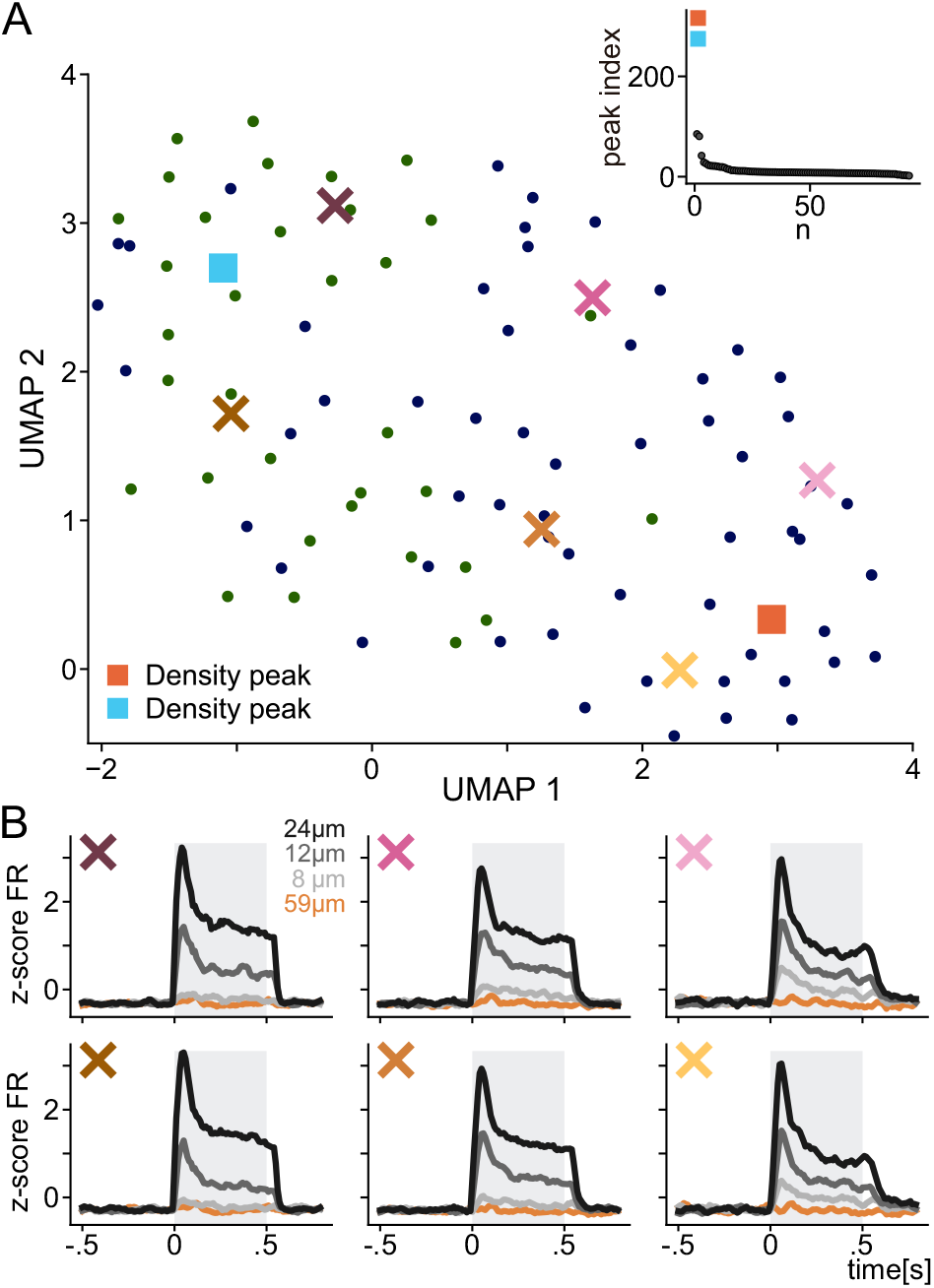
(A) 2-D non-linear projection of the normalized neural activity using UMAP. Neurons from area 3b are represented with green circles while neurons from area 1 with blue circles. Orange and blue squares indicate the two highest density peaks. Inset plot shows local density peaks from the highest to the lowest. Colored crosses indicate the center of the weighted local averages of firing rate profiles. (B) Transformation of the activity profiles. Colored traces represent the weighted averages of firing rate profiles for the supraliminal (24 μm), liminal (12 μm) and subliminal (8 μm) ranges centered on each X mark. Acoustic profiles for the supraliminal condition (59 dB) are shown in orange.

